# Spc1 regulates substrate selection for signal peptidase

**DOI:** 10.1101/2021.02.02.429376

**Authors:** Chewon Yim, Yeonji Chung, Jeesoo Kim, IngMarie Nilsson, Jong-Seo Kim, Hyun Kim

## Abstract

Signal peptidase (SPase) cleaves the signal sequences (SSs) of secretory precursors. It contains an evolutionarily conserved membrane protein subunit, Spc1 that is dispensable for the catalytic activity of SPase, and its role remains elusive. In the present study, we investigated the function of yeast Spc1. First, we set up an *in vivo* SPase cleavage assay using secretory protein CPY variants with SSs modified in the *n* and *h* regions. When comparing the SS cleavage efficiencies of these variants in cells with or without Spc1, we found that signal-anchored sequences become more susceptible to cleavage by SPase without Spc1. Further, SPase-mediated processing of transmembrane (TM) segments in model membrane proteins was reduced upon overexpression of Spc1. Spc1 was co-immunoprecipitated with membrane proteins carrying uncleaved signal-anchored or TM segments. These results suggest a role of Spc1 in shielding TM segments from SPase action, thereby contributing to accurate substrate selection for SPase.

## Introduction

Signal peptidase (SPase) is an evolutionarily conserved protease that cleaves signal sequences (SSs) of secretory precursors targeted to the plasma membrane in prokaryotes or the endoplasmic reticulum (ER) in eukaryotes. Processing occurs co- or post-translationally when a nascent chain passes through the Sec translocon (Lyko et al., 1995; Wollenberg and Simon, 2004).

In prokaryotes, SPase I (leader peptidase) functions as a monomer, whereas eukaryotic SPase is a heterooligomer consisting of membrane protein subunits, all of which are conserved from yeast to humans (Spc1/SPCS1, Spc2/SPCS2, Spc3/SPCS3 and Sec11/SEC11A and Sec11/SEC11C) (Antonin et al., 2000; Dalbey et al., 2017; Dalbey and Von Heijne, 1992; Evans et al., 1986; Fang et al., 1996; Greenburg et al., 1989; Shelness and Blobel, 1990; YaDeau et al., 1991; Zwizinski and Wickner, 1980).

Sec11 and Spc3 are required for the catalytic activity of eukaryotic SPase and essential for cell viability in yeast. Both Sec11 and Spc3 are single-pass membrane proteins with the C-terminal domain facing the lumen, and their luminal domains exhibit sequence homology to the leader peptidase active domain (Fang et al., 1997; VanValkenburgh et al., 1999). Spc2 was found to associate with a beta subunit of the Sec61 complex in both yeast and mammals, suggesting its role in an interaction between the SPase complex and the Sec61 translocon (Antonin et al., 2000; Kalies et al., 1998).

Spc1 was first identified from a homology search of mammalian Spc12 (SPCS1) and genetic interaction with Sec11 in yeast (Fang et al., 1996). While Spc1 is dispensable for cell viability in yeast, deletion of SPC12 (Spc1 homolog) in *Drosophila* causes a developmental defect, indicating its crucial role in higher eukaryotes (Haase Gilbert et al., 2013). In the yeast strain lacking Spc1, signal peptides of secretory precursors were efficiently cleaved (Fang et al., 1996; Mullins et al., 1996); hence, the role of Spc1 in SPase remains puzzling.

SSs have common characteristics: a hydrophobic core containing consecutive nonpolar amino acids that can form at least two turns in an α-helix (*h* region) flanked by N-terminal (*n* region) and C-terminal (*c* region) polar and charged residues (von Heijne, 1985). While a tripartite structure is found in all SSs, the overall and relative lengths of the *n, h*, and *c* regions, the hydrophobicity of the *h* region, and the distribution of charged residues in the *n* and *c* regions greatly vary among them, making SSs uniquely diverse (Choo and Ranganathan, 2008; Choo et al., 2008; Gierasch, 1989; Hegde and Bernstein, 2006; von Heijne, 1985).

SPase recognizes a cleavage motif that includes small, neutral amino acids at the −3 and −1 positions relative to the cleavage site (Bird et al., 1990; Dalbey and Von Heijne, 1992; von Heijne, 1990). The structure of bacterial SPase shows binding pockets for small residues in the active site (Paetzel et al., 1998). However, not all SSs with an optimal cleavage site are processed by SPase (Nilsson et al., 1994; Yim et al., 2018). On the other hand, a signal anchor of sucrase-isomaltase was found to be cleaved when a single amino acid was substituted within a signal anchor sequence, illustrating that subtle changes in and around the TM domain can make it a substrate for SPase (Hegner et al., 1992). These observations imply that SPase recognizes certain characteristics in SSs in addition to the cleavage site, yet it remains unknown how SPase discriminates signal-anchored or TM segments of membrane proteins while accurately selecting signal peptides (SPs) of secretory precursors.

To investigate the process by which SPase selects substrates, we first set up an *in vivo* SPase cleavage assay in *Saccharomyces cerevisiae* using carboxypeptidase Y (CPY) variants carrying SSs of systematically varied length and hydrophobicity. With this approach, we defined the substrate range of SPase in terms of *n* and *h* region features of SSs in yeast. Next, we undertook to explore the role of Spc1. We assessed and compared the SS cleavage efficiencies of the CPY variants in cells with or without Spc1. We observed that membrane-anchored, internal SSs were more efficiently cleaved in the absence of Spc1. Mutagenesis analysis at the cleavage site showed that recognition and usage of cleavage sites by Spase was unaffected with or without Spc1. Further, cleavage of a TM segment in model membrane proteins was enhanced in the absence of Spc1 but reduced upon overexpression of Spc1. Collectively, our data show that SPase selects substrates based on the *n* and *h* regions of the signal sequence and becomes more prone to include signal-anchored and TM domains for processing without Spc1. These results suggest that Spc1 protects TM segments of membrane proteins from being cleaved by SPase, implicating its role in regulating the substrate selection for SPase.

## Results

### Defining the substrate spectrum of SPase in S. cerevisiae

Previously, we observed that a secretory protein, carboxypeptidase Y (CPY), having a hydrophobic SS (CPY(*h*); *h* for high hydrophobicity) and the same precursor with the N-terminal extension localized differently (N26CPY(*h*)) (Yim et al., 2018). The former was found in the soluble fraction and migrated faster, while the latter was found in membrane pellets upon carbonate extraction and migrated more slowly (Fig. S1A). These data suggested that SS cleavage may differ depending on the length of the N-terminus preceding the SS (N-length), and we undertook to determine the relationship between the N-length and efficiency of SS cleavage by systematic truncation of the N-terminus of N26CPY (N26CPYt(*h*))(Fig. 1A and Table 1).

**Figure 1.**
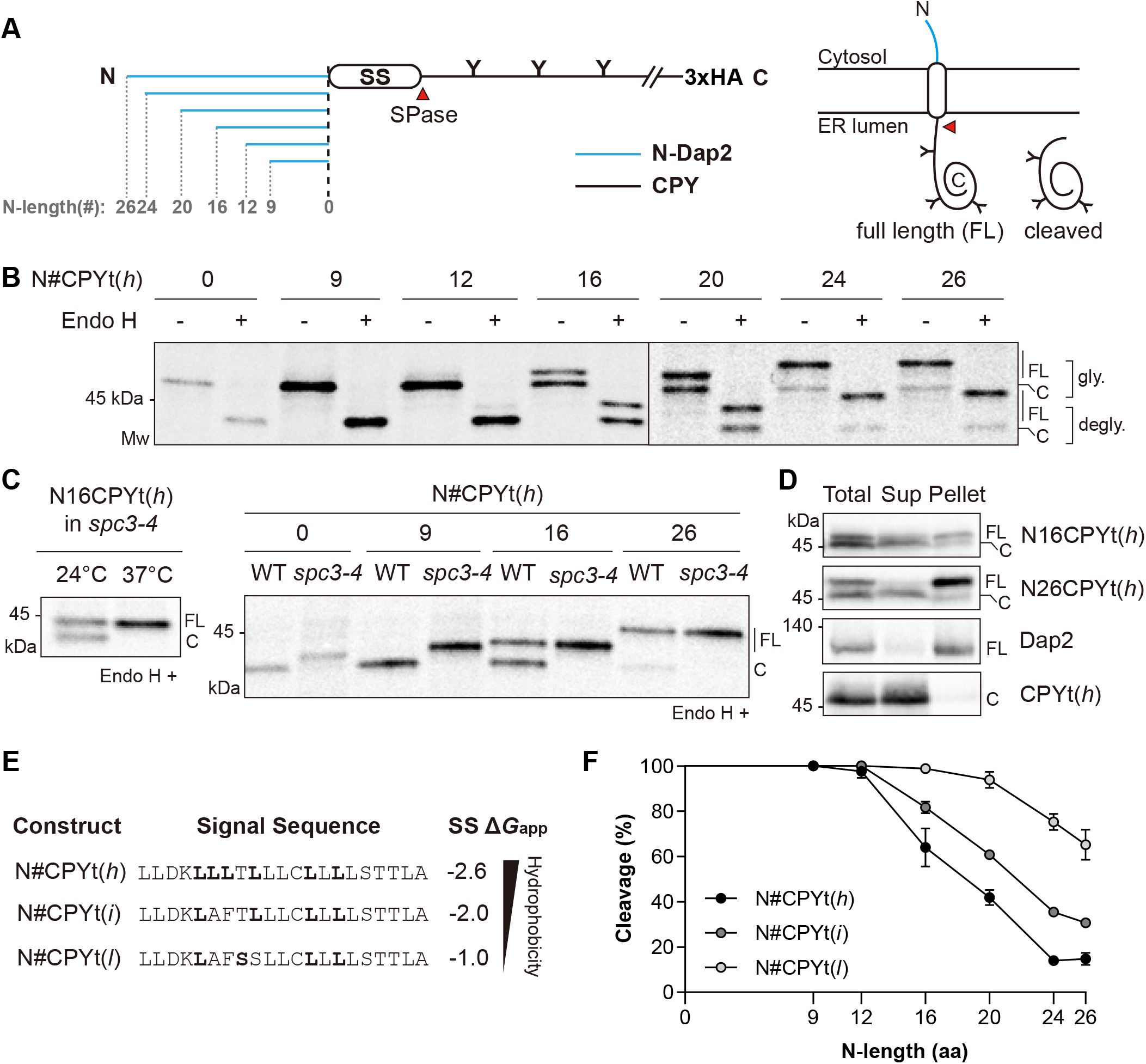
Signal sequence (SS) processing by SPase depends on the *n* region length and *h* region hydrophobicity of SSs. (A) Schematics of N#CPYt constructs. *Left*, Blue lines indicate N-terminal extension derived from the N-terminus of the yeast membrane protein Dap2 (N-Dap2), and black line indicates the yeast vacuole protein CPY. Numbers indicate the extended amino acids. N-linked glycosylation sites are shown as Y. SS, signal sequence; HA, hemagglutinin tag. *Right*, a cartoon of possible forms of N#CPYt in the ER. (B) Yeast transformants carrying the indicated N#CPYt(*h*) constructs were radiolabeled for 5 min at 30°C, immunoprecipitated with an anti-HA antibody, subjected to SDS-PAGE and analyzed by autoradiography. Endoglycosidase H (Endo H) treatment was performed prior to SDS-PAGE. FL, full length; C, cleaved; gly.; glycosylated speicies; degly.;deglycosylated species. (C) The indicated N#CPYt(*h*) variants in the WT or *spc3-4* strain were analyzed as in (B), except that N#CPYt(*h*) variants in the *spc3-4* strain were incubated at 37°C for 30 min prior to radiolabeling and radiolabeled at 37°C. (D) The indicated CPY variants and Dap2 were subjected to carbonate extraction, and the resulting protein samples were detected by Western blotting using an anti-HA antibody. (E) Hydrophobicities of the N#CPYt variant SSs were predicted by the ΔG predictor (ΔG_app_ (kcal/mol)) (http://dgpred.cbr.su.se/). (F) The relative amounts of SPase-processed species over glycosylated products for each CPY variant were measured and plotted (cleavage (%)). The x-axis indicates the number of amino acids preceding the SS (N-length). At least three independent experiments were carried out, and the average is shown with the standard deviation.

**Table 1.**
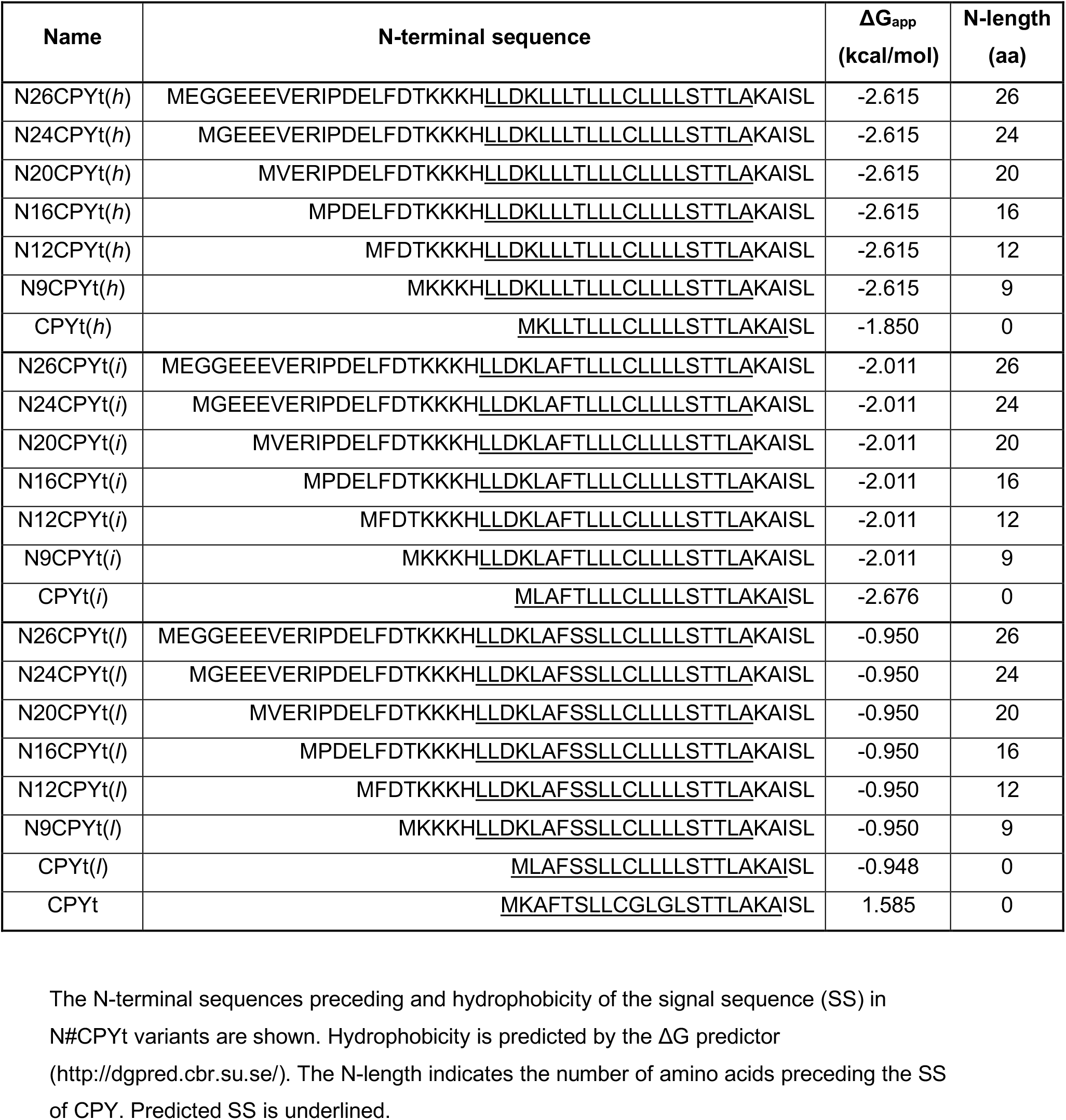
List of CPY variants used in this study.

For better separation of the size of a cleaved and an uncleaved species on SDS-PAGE, the C-terminus was shortened to residue 323 of CPY. To capture the early stage of protein translocation and processing, yeast cells carrying N#CPYt(*h*) variants (N# denotes the number of amino acids extended at the N-terminus) were radiolabeled with [^35^S]-Met for 5 min. Radiolabeled proteins were immunoprecipitated with anti-HA antibodies directed to the HA epitope at their C-terminus, subjected to SDS-PAGE and analyzed by autoradiography. Proper targeting and translocation of CPY to the ER was determined by assessing the glycosylation status of CPY, as it contains three N-linked glycosylation sites, which take place in the ER lumen. All N#CPYt(*h*) variants were sensitive to treatment with endoglycosidase H (Endo H), which removes N-linked glycans, indicating that they were efficiently translocated into the lumen (Fig. 1B).

Two bands were detected for the longer N-length variants (N16CPYt(*h*) to N26CPYt(*h*)) even with Endo H treatment, indicating that they are proteins of two different sizes in the ER (Fig. 1B). When the size of CPYt(*h*) and N16CPYt(*h*) was compared to that of CPYt(wt) (CPYt possessing the original SS), CPYt(*h*) and the smaller size form of N16CPYt(*h*) migrated the same as CPYt(wt), SS of which is efficiently cleaved by SPase. Thus, CPYt(*h*) is fully cleaved and N16CPYt(*h*) is partially cleaved by SPase (Fig. S1B). We also prepared an N-terminal SS-deleted version of CPYt(*h*) (mCPYt), which was expressed *in vitro*, and compared its size with the Endo H-treated sample of N16CPYt(*h*) expressed *in vivo*. A fast-migrated, deglycosylated band of N16CPYt(*h*) and an *in vitro* translated product of mCPYt were resolved at the same size on an SDS-gel, confirming that the former is an SS-cleaved CPY (Fig. S1C).

To further confirm the SPase-mediated cleavage, selective N#CPYt(*h*) variants were expressed in the *spc3-4* strain, which exhibits a temperature-sensitive defect in SPase activity (Fang et al., 1997). When N16CPYt(*h*) was radiolabeled at a permissive temperature of 24°C, two forms appeared, whereas a lower band was not observed in cells radiolabeled at the nonpermissive temperature of 37°C, indicating that the lower band resulted from SPase activity (Fig. 1C). CPYt(*h*) and N9CPYt(*h*) variants expressed at 37°C in the *spc3-4* strain migrated slower than those expressed in the wild-type (WT) strain, and fast-migrated bands of the N16CPYt(*h*) and N26CPYt(*h*) variants in the *spc3-4* strain were no longer detected when labeled at 37°C (Fig. 1C).

Finally, we determined the localization of SS-cleaved and -uncleaved species by carbonate extraction followed by Western blotting. SS-cleaved forms of N16CPYt(*h*) and N26CPYt(*h*) variants were found in soluble fractions, while the uncleaved forms were mainly found in pellet fractions, indicating that the latter became membrane-anchored (Fig. 1D).

These data show that SSs of N#CPYt(*h*) variants with shorter N-lengths (CPYt(*h*), N9CPYt(*h*) and N12CPYt(*h*)) were efficiently cleaved, whereas cleavage gradually decreased for variants with longer N-lengths, indicating that SSs with shorter N-lengths are better substrates for SPase than SSs with longer N-lengths.

We next investigated the effect of the hydrophobicity of SS on cleavage by SPase. Sets of N#CPYt variants having SSs of intermediate hydrophobicity (N#CPYt(*i*), *i* for intermediate) and low hydrophobicity (N#CPYt(*l*), *l* for low) were prepared and analyzed by 5 min pulse labeling as above (Figs. 1E and S1E). The relative amounts of SS-cleaved species among the glycosylated products were quantified (% cleavage) (Fig. 1F). The SS cleavage profiles of the (N#CPYt(*i*)) and N#CPYt(*l*) variants were shifted to the right, and the estimated threshold N-length (50% SS cleavage by applying a trend line on the graphs) increased as the SS became less hydrophobic (∼19 for CPYt(*h*), ∼22 for CPYt(*i*), and >26 for CPYt(*l*)) (Fig. 1F).

These data show that the N-length and hydrophobicity of SSs are two critical determinants based on which SPase distinguishes substrates (cleavable SSs, SPs) from nonsubstrates (uncleavable SSs, TMs).

### Internal SSs are more efficiently cleaved by SPase lacking Spc1

The eukaryotic SPase has multiple subunits and the functions of each subunit remain poorly defined. We set out to investigate the role of Spc1, a small membrane protein subunit (Fang et al., 1996; Kalies and Hartmann, 1996). Spc1 spans the ER membrane twice, with both termini facing the cytoplasm with a very short loop in the lumen (Fig. 2A).

**Figure 2.**
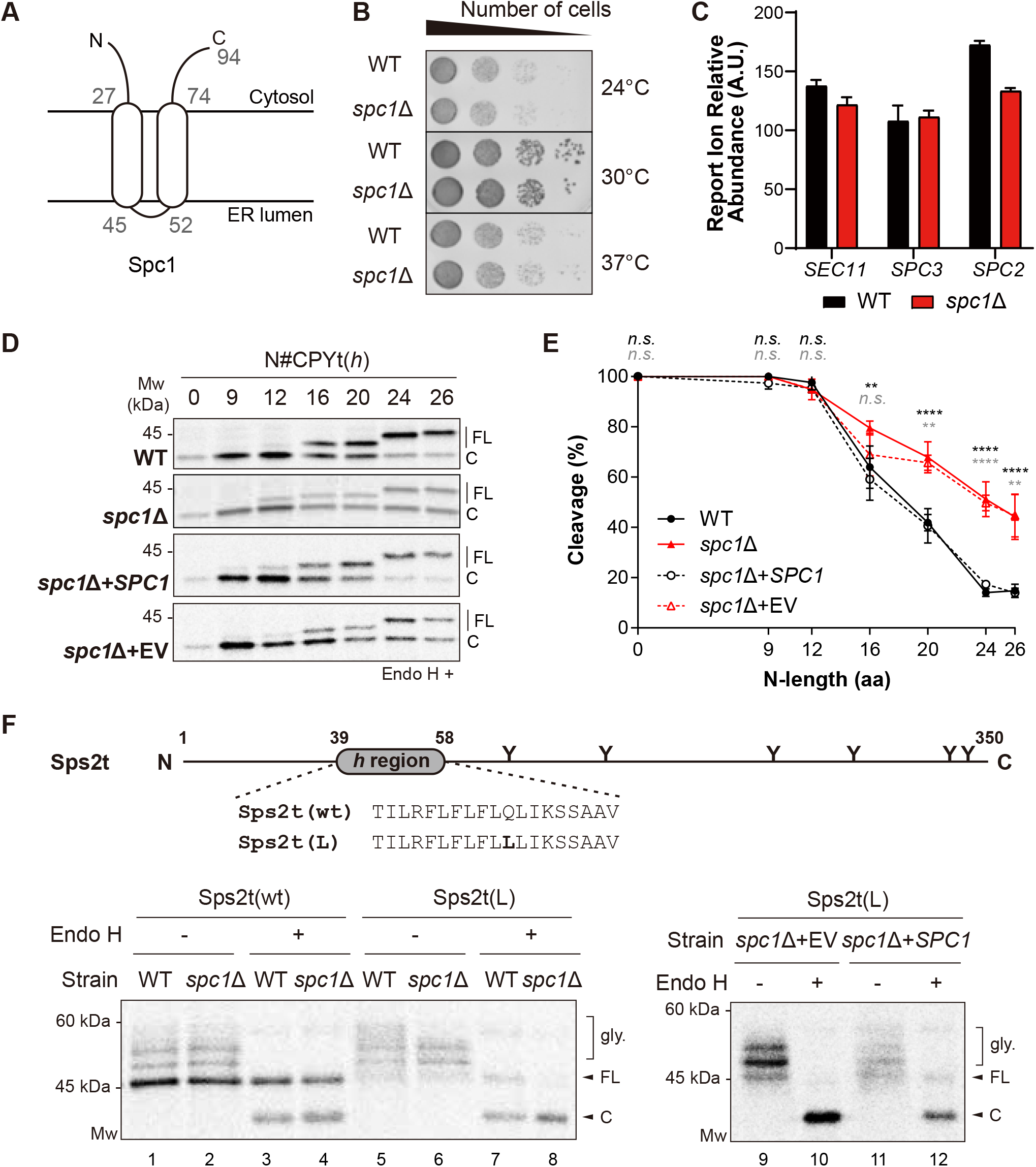
Cleavage of internal SSs is enhanced in the absence of Spc1. (A) Schematics of the membrane topology of Spc1. Numbers indicate the ends of two TM domains of Spc1. (B) WT and *spc1*Δ cells were serially diluted from 0.2 OD_600_ cells and grown on YPD medium for 1 day at the indicated temperatures. (C) The abundance of other SPC subunits, Sec11, Spc3 and Spc2, in the WT or *spc1*Δ strain was assessed by mass spectrometry analysis. The standard deviation of three repeats are shown. (D) N#CPYt(*h*) variants in the indicated strains were assessed as in Fig.1B. (E) Cleavage (%) of N#CPYt(*h*) variants in the WT, *spc1*Δ, *spc1*Δ+*SPC1*, and *spc1*Δ+EV strains were analyzed. At least three independent experiments were carried out and the average is shown with the standard deviation. p-values between WT and *spc1*Δ and between *spc1*Δ+EV and *spc1*Δ+*SPC1* were calculated by multiple *t*-tests and shown in black and grey colors, respectively; *n*.*s*., p>0.05; **, p≤0.01; ****, p≤0.0001. (F) Schematics (*top*) and processing of Sps2t variants in WT, *spc1*Δ, *spc1*Δ+*SPC1*, and *spc1*Δ+EV strains (*bottom*). Experimental procedures were carried out as in Fig. 1B. FL, full length; C, cleaved.

First, we prepared a *spc1*Δ strain and assessed its growth phenotype. No growth defect was observed at any tested temperature, as seen in an earlier study (Fig. 2B) (Fang et al., 1996). To check if the deletion of Spc1 affects the stability of the other subunits in SPase, we carried out mass spectrometry analysis to compare the abundance of Sec11, Spc3 and Spc2 in WT and *spc1*Δ cells. Although the abundance of the nonessential subunit Spc2 was slightly reduced in the *spc1*Δ strain, the abundance of Sec11 and Spc3, which are the catalytic components of SPase, was unchanged (Fig. 2C), indicating that Spc1 deletion hardly affects the stability of other subunits in the complex. Further, SS cleavage of shorter N#CPYt variants in *spc1*Δ cells occurred efficiently, indicating that SPase activity is not impaired in the absence of Spc1 (Figs. 2D, E and S2).

For N#CPYt(*h*) variants with N-lengths longer than 16 in *spc1*Δ cells, SS cleavage was increased compared to that in WT cells (Figs. 2D and E). The difference in the SS cleavage efficiency in *spc1*Δ and WT cells became larger as the N-length became longer (Figs. 2D and E). SS cleavage was also assessed in a *spc1*Δ strain carrying a plasmid with *SPC1* under its own promoter or an empty vector (EV). Cleavage efficiencies of N#CPYt(*h*) variants in a *spc1*Δ strain with *SPC1* were restored to the level in the WT strain, confirming that increased cleavage of longer N-length variants in the *spc1*Δ strain is due to the absence of Spc1 (Figs. 2D and E).

Less hydrophobic N#CPYt(*i*) and N#CPYt(*l*) sets showed similar cleavage patterns: cleavage efficiency increased for the longer N-length variants when Spc1 was absent and restored when *SPC1* was re-expressed in the *spc1*Δ strain (Figs. S2A and B). Thus, these data show that SPase lacking Spc1 becomes more prone to cleave membrane-anchored, internal SSs.

### Processing of Sps2

We wondered whether Spc1-regulated processing of internal SSs also occurs in natural proteins and searched for yeast endogenous proteins possessing an internal SS. We found Sps2, a protein involved in sporulation and localized to the plasma membrane and cell wall in *S. cerevisiae* (Coluccio et al., 2004)(Fig. 2F).

To facilitate the separation of SS-cleaved and -uncleaved forms on SDS-gels, C-terminus-truncated Sps2 (Δ351-502, Sps2t) was used for the cleavage assay (Fig. 2F). After 5 min of radiolabeling, the unglycosylated protein was detected in WT and *spc1*Δ strains, indicating inefficient targeting to the ER (Fig. 2F, lanes 1-4). Since inefficient ER targeting obscures analysis of cleavage, a single amino acid substitution in the *h* region was made to improve ER targeting (Sps2t(L))(Fig. 2F, lanes 7-8). Sps2t(L) was first expressed in the *spc3-4* strain and radiolabeled at 37°C, the nonpermissive temperature to analyze SPase-mediated cleavage. However, untargeted proteins accumulated at 37°C, thus we assessed the cleavage at 33°C, a semipermissive temperature without compromising ER targeting and confirmed that the cleaved product of Sps2t(L) is generated by SPase (Fig. S2C). Sps2t(L) was expressed in WT and *spc1*Δ strains, and when the cleaved product was assessed upon Endo H digestion, only the SS cleaved form was detected in *spc1*Δ cells whereas the full-length form was detected in WT cells (Fig. 2F, lanes 7-8). Sps2t(L) was also expressed in the *spc1*Δ strain with an EV or *SPC1*, and the data confirmed that the full-length protein was more readily cleaved in the absence of Spc1 (Fig. 2F, lanes 10 and 12). These results show that cleavage of the internal SS of Sps2 can also be regulated by Spc1.

### Recognition and usage of the SS cleavage site by SPase is unchanged without Spc1

We wondered whether the expanded substrate range of SPase in *spc1*Δ cells is due to altered recognition and usage of cleavage sites by SPase lacking Spc1 and investigated the cleavage sites of CPY variants in WT and *spc1*Δ cells.

When the SS cleavage sites of CPY variants were searched with SignalP-5.0 (http://www.cbs.dtu.dk/services/SignalP/) (Almagro Armenteros et al., 2019), two sites were predicted for all CPYt variants (including wild-type CPY) with equal probabilities (0.492 for the upstream cleavage site and 0.482 for the downstream cleavage site, Figs. 3A and S3A). We referred to the upstream and downstream cleavage sites as cleavage sites 1 and 2, respectively, and denoted residues around cleavage site 2 with ‘(*e*.*g*., −3’, −1’, Fig. 3A).

**Figure 3.**
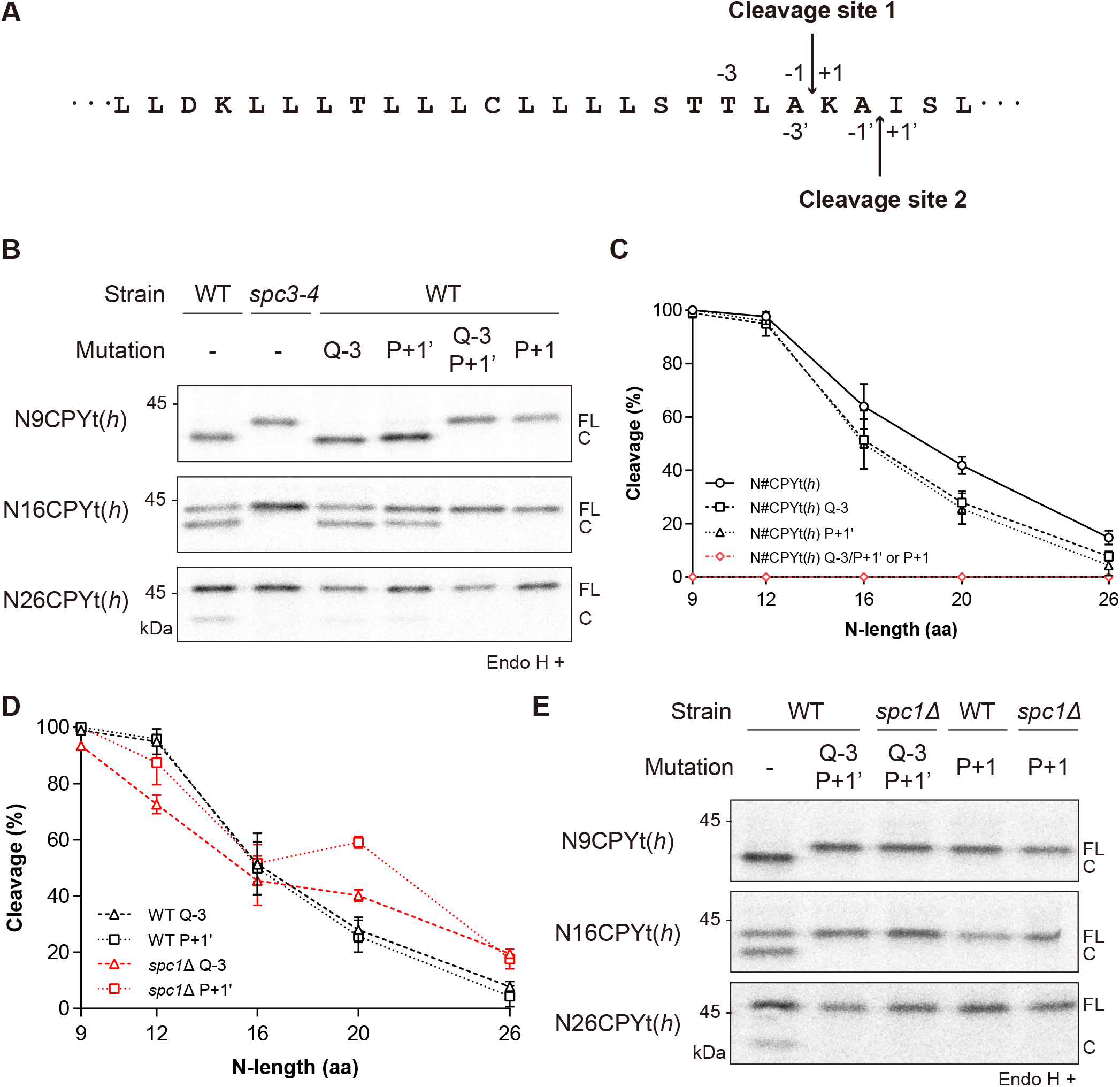
Recognition and usage of the SS cleavage site by SPase is unchanged in the *spc1*Δ strain. (A) Two cleavage sites are present in the SS of N#CPYt(*h*): cleavage site 1 and cleavage site 2 are indicated as downward and upward arrows, respectively. (B) The indicated cleavage site mutants of N#CPYt(*h*) variants in the WT or *spc3-4* strain were radiolabeled for 5 min at 30°C (37°C for *spc3-4*), immunoprecipitated by anti-HA antibodies, subjected to SDS-PAGE and Endo H treatment and analyzed by autoradiography. (C) Cleavage (%) of the cleavage site mutants in (B) is analyzed as in Fig. 1F and compared. (D) Cleavage (%) of N#CPYt(*h*) variants with Q-3 or P+1’ mutations in the WT or *spc1*Δ strain is compared. At least three independent experiments were carried out, and the average is shown with the standard deviation. (E) The indicated N#CPYt(*h*) variants lacking canonical cleavage sites in the WT or *spc1*Δ strain were radiolabeled. FL, full length; C, cleaved.

To identify which cleavage site is used, cleavage site 1 or 2 was selectively eliminated one at a time by single amino acid substitution in N#CPYt(*h*) variants. Given that the canonical SS cleavage sites follow the −3, −1 rule and proline (P) at the +1 position with respect to the cleavage site inhibits SS processing (Cui et al., 2015; Nilsson and von Heijne, 1992) (Barkocy-Gallagher and Bassford, 1992), a residue at the −3 position in cleavage site 1 was replaced with the polar and bulky residue glutamine (Q-3), and the +1’ position in cleavage site 2 was replaced with proline (P+1’) (Fig. 3B). Prediction by SignalP 5.0 showed a single cleavage site for each mutant, indicating that Q-3 and P+1’ substitutions disrupt cleavage sites 1 and 2, respectively (Fig. S3). To confirm that cleavage site 1 or 2 is selectively eliminated by Q-3 or P+1’ substitution, N#CPYt(*h*) variants carrying the double mutation Q-3/P+1’ were also prepared.

Cleavage of N#CPYt(*h*) variants possessing cleavage site mutations was assessed by 5 min pulse experiments, and the data were compared with the cleavage profile of N#CPYt(*h*) variants. Although a slightly decreased cleavage of N16CPYt(*h*), N20CPYt(*h*) and N26CPYt(*h*) was observed upon inhibition at cleavage site 1 (Q-3) or site 2 (P+1’), the overall pattern of the cleavage profile remained the same regardless of whether they contained both cleavage sites or only cleavage site 1 or 2, suggesting that SPase recognizes and uses both sites efficiently (Figs. 3B and C). On the other hand, double mutation Q-3/P+1’ completely abolished SS processing of all variants. When proline was introduced at the +1 position for cleavage site 1 (P+1), which is the −2’ position for cleavage site 2, SS cleavage was also completely blocked, indicating that both sites were inhibited by the presence of proline at this position (Figs. 3B and C). We wondered whether N9CPYt(*h*), a short N-length variant, was present in the membrane when uncleaved and carried out carbonate extraction. N9CPYt(*h*) with the Q-3/P+1’ mutation was found in the membrane pellet, showing that the protein becomes membrane-anchored when unprocessed by SPase (Fig. S3D).

To determine if SPase lacking Spc1 uses different cleavage sites for processing, we analyzed SS processing of the cleavage site variants in *spc1*Δ cells. Cleavage of N20CPYt(*h*) and N26CPYt(*h*) variants with Q-3 or P+1’ mutations in *spc1*Δ cells increased compared to that in WT cells (Fig. 3D). These results indicate that recognition and usage of cleavage sites for SPase without Spc1 are unchanged. Next, we set out to determine whether SPase lacking Spc1 uses a noncanonical SS cleavage site, thereby evading the −3, −1 rule for processing. Three N#CPYt(*h*) variants with the Q-3/P+1’ or P+1 mutation that eliminated both canonical SS cleavage sites were expressed in *spc1*Δ cells, and their cleavage was assessed (Fig. 3E). As in WT cells, no cleavage was detected for these sets of variants, indicating that SPase still processes the canonical SS cleavage sites, even in the absence of Spc1 (Fig. 3E).

### SPase-mediated cleavage of TM segments in membrane proteins is enhanced in the absence of Spc1

Observing that SPase lacking Spc1 includes internal, membrane-anchored SSs for processing, we reasoned that TM segments of membrane proteins may also be subjected to SPase-mediated cleavage in *spc1*Δ cells. To test this idea, LepCC, *E. coli* leader peptidase (Lep)-derived membrane proteins, were used.

LepCC proteins contain the engineered TM2 composed of Leu residues followed by a cleavage cassette (VPSAQA↓A, ↓ is the cleavage site of SPase, Fig. S4A) (Nilsson et al., 1994). An earlier study showed that SPase-mediated cleavage after TM2 of these proteins was dependent on the number of Leu residues in TM2 when determined *in vitro* with dog pancreatic microsomes; TM2 variants with a shorter stretch of Leu residues were cleaved by SPase, while TM2 variants with a longer stretch of Leu residues were uncleaved (Nilsson et al., 1994).

We deleted the N-terminus including the first TM of LepCC, to generate signal-anchored LepCC versions with 14 leucines (LepCCt(14L)), 17 leucines (LepCCt(17L)), and 20 leucines (LepCCt(20L)) in their TMs and subcloned the fragments in a yeast expression vector (LepCCt) (Fig. 4A). All three constructs were expressed in yeast cells. Upon Endo H treatment, the band shifted down for all LepCCt variants, indicating efficient translocation and membrane insertion in the yeast ER (Fig. 4B). For LepCCt(14L), the size of the major band was smaller than the expected full-length protein (Fig. 4B). To determine whether smaller band size resulted from SPase-mediated processing, we adopted two strategies. First, an SS cleavage site was destroyed by introducing proline at the +1 position in LepCCt(14L); second, LepCCt(14L) was radiolabeled in the *spc3-4* strain at the nonpermissive temperature (Fig. 4C). A slowly migrated full-length band was detected in LepCCt(14L) with a cleavage site mutation (P+1) or when expressed in the *spc3-4* strain at the nonpermissive temperature, whereas a fast-migrated product was predominant when expressed in the WT strain, confirming that LepCCt(14L) was processed by SPase in yeast. Albeit less prominent, LepCCt(17L) also generated a fast-migrated band, indicating that it is a substrate for SPase as well (Fig. 4B).

**Figure 4.**
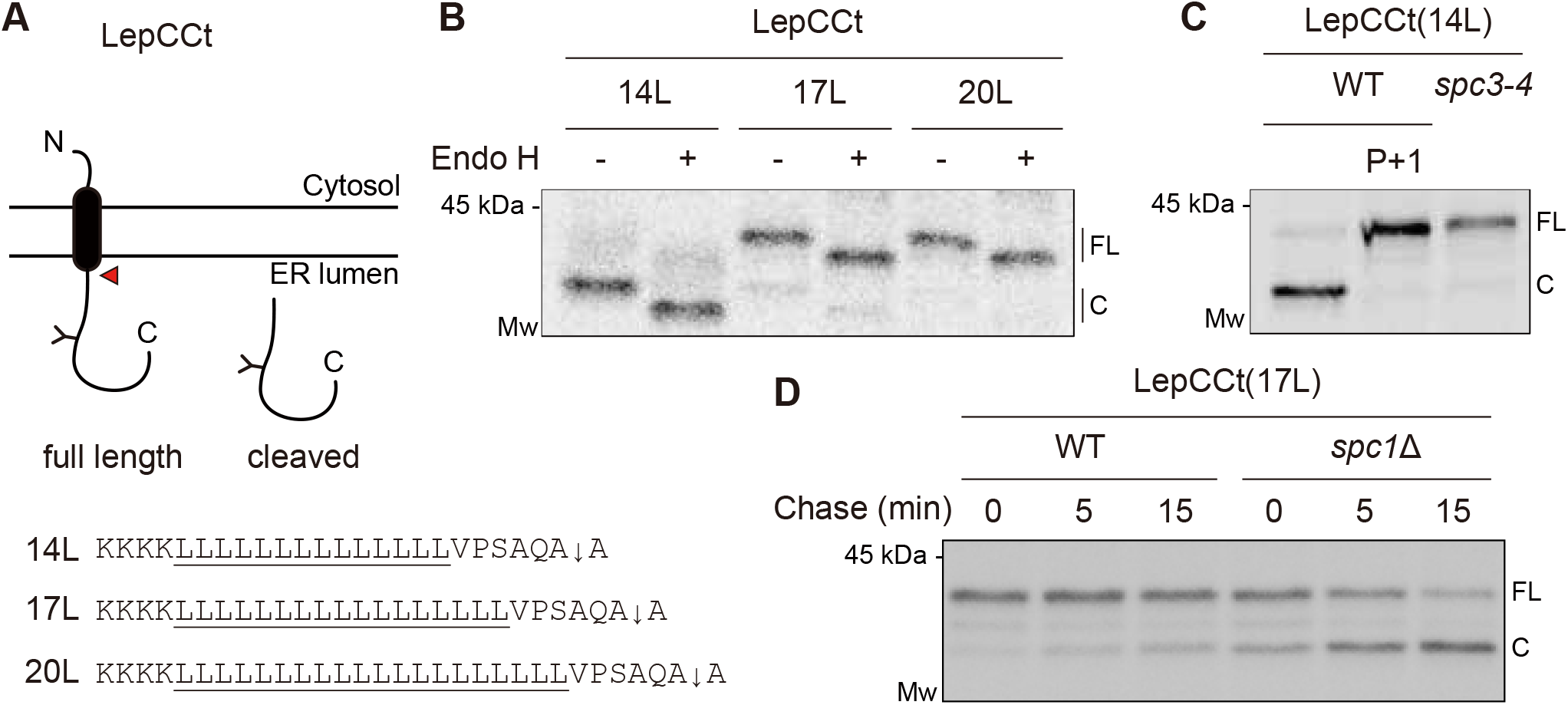
SPase-mediated processing of signal-anchored proteins is enhanced in the *spc1*Δ strain. (A) Schematics of LepCCt. The TM domain is colored black, and an N-linked glycosylation site is indicated as Y. Flanking and TM sequences including the cleavage site (↓) are shown for three LepCCt variants. A red arrowhead points to cleavage by SPase. (B) The indicated LepCCt variants in WT cells were radiolabeled for 5 min at 30°C and subjected to immunoprecipitation for SDS-PAGE and autoradiography. Protein samples were treated with or without Endo H prior to SDS-PAGE. FL, full length; cleaved, cleaved species. (C) The LepCCt(14L) construct in the WT or *spc3-4* strain was radiolabeled and analyzed as in (B). P+1 in LepCCt(14L) indicates proline substitution in the +1 position relative to the cleavage site. (D) LepCCt(17L) in the WT or *spc1*Δ strain was radiolabeled for 5 min, chased for the indicated time points at 30°C, immunoprecipitated with anti-HA antibodies, subjected to SDS-PAGE, and analyzed by autoradiography.

We next traced processing of LepCCt(17L) variant in the WT or *spc1*Δ strain by pulse-chase experiments (Fig. 4D). A cleaved product of LepCCt(17L) in the *spc1*Δ strain significantly increased at 0 min compared to that expressed in the WT strain and further increased in the following chase time, indicating that SPase-mediated cleavage continued post-translationally (Fig. 4D). We also determined cleavage of double-spanning LepCC variants in WT and *spc1*Δ cells and observed that SPase-mediated cleavage of a TM domain increased in *spc1*Δ cells, similar to cleavage of single-spanning LepCCt variants (Figs. S4A and B). These data suggest that longer TM segments normally evade SPase-mediated processing, but they are subjected to cleavage by SPase when Spc1 is absent.

Next, to determine the effect of the hydrophobicity of TM segments on SPase-mediated cleavage, we tested another set of *E. coli* Lep-derived membrane proteins harboring the engineered TM2 composed of Leu and Ala residues with a fixed length of 19 residues (LepH2, Fig. 5A) (Lundin et al., 2008). The segment becomes more hydrophobic with an increasing number of Leu residues. Previously, it was shown that LepH2 variants undergo SPase-mediated cleavage *in vitro* in dog pancreas microsomes and *in vivo* in yeast cells (Lundin et al., 2008). We confirmed that the cleaved fragment was generated by SPase by expressing LepH2(3L) in the *spc3-4* strain at the nonpermissive temperature (Fig. 5B).

**Figure 5.**
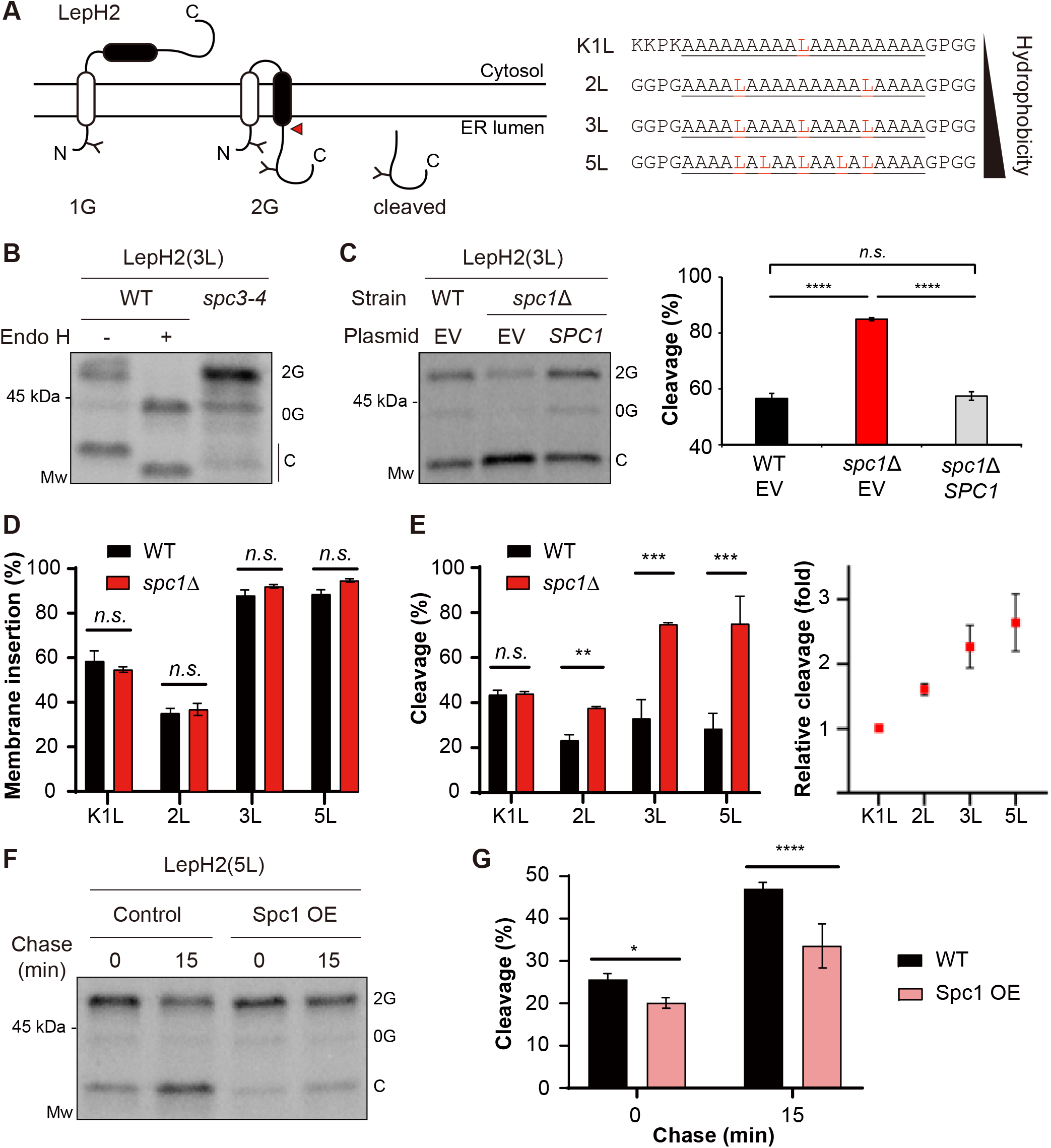
SPase-mediated processing of double-spanning membrane proteins is modulated by Spc1. (A) Schematics of LepH2. The second hydrophobic (H) segment of varying hydrophobicity is colored black. Amino acid sequences of the H segment are shown and underlined, with N- and C-terminal flacking residues. N-linked glycosylation sites are indicated as Y. A red arrowhead points to cleavage by SPase. (B) LepH2(3L) in the WT or *spc3-4* strain was radiolabeled for 5 min at 30°C. WT samples were treated with Endo H prior to SDS-PAGE. 0G, nonglycosylated form; 2G, doubly glycosylated form; C, cleaved form after membrane insertion. (C) LepH2(3L) in the WT+EV, *spc1*Δ+EV or *spc1*Δ+*SPC1* strain was radiolabeled for 5 min at 30°C and analyzed. *Left*, autoradiogram of a representative blot is shown. *Right*, cleavage (%) of the LepH2 H segments was calculated as cleaved/(2G+cleaved)*100 (%) from three independent experimental measurements. p-values were calculated by multiple *t*-tests; *n*.*s*., p>0.05; ****, p≤0.0001. (D) Membrane insertion of the H segment in LepH2 variants was measured as doubly glycosylated products/(total-0G)*100 (%). (E) *Left*, cleavage (%) of the LepH2 H segments was calculated as in (C). *Right*, relative cleavage of the LepH2 H segments in *spc1*Δ cells compared to that in WT cells was calculated as cleavage (%) in *spc1*Δ cells/cleavage (%) in WT cells and plotted for each LepH2 variant. (F) LepH2(5L) in WT cells harboring control vector or Spc1 overexpression (OE) vector. Transformants were subjected to radiolabeling for 5 min at 30°C followed by chasing for the indicated time points. (G) Cleavage (%) of LepH2(5L) in (F) was calculated as in (C). For all the experimental sets, at least three independent experiments were carried out, and the average is shown with the standard deviation. p-values were calculated by multiple *t*-tests; *n*.*s*., p>0.05; ****, p≤0.0001.

LepH2(3L) was expressed in WT and *spc1*Δ strains carrying an EV or a plasmid bearing the *SPC1* gene to assess whether re-expression of Spc1 restores LepH2(3L) processing as in the WT strain. Indeed, cleavage of LepH2(3L) in a *spc1*Δ strain with *SPC1* was comparable to that in the WT strain with an empty vector, reaching ∼55%, whereas cleavage of LepH2(3L) in the *spc1*Δ strain with an EV was resulted in ∼85%. These data show that Spc1 regulates SPase action on processing of LepH2 (Fig. 5C).

Additional LepH2 variants of varying hydrophobicities were expressed in WT and *spc1*Δ strains and analyzed. Membrane insertion of TM2 of LepH2 variants in WT and *spc1*Δ cells remained unchanged, demonstrating that deletion of Spc1 does not interfere with membrane insertion of TM2 (Fig. 5D). However, cleavage of LepH2 constructs in the *spc1*Δ strain increased in a hydrophobicity-dependent manner; cleavage of hydrophobic LepH2(3L) and LepH2(5L) was significantly increased in the *spc1*Δ strain (>2-fold) (Fig. 5E). LepH2(K1L) contains positively charged N-terminal flanking residues that enhance membrane insertion of TM2 despite low hydrophobicity. Although membrane insertion of LepH2(K1L) was better than that of LepH2(2L), cleavage of LepH2(K1L) in *spc1*Δ cells did not increase whereas cleavage of LepH2(2L) did. These data collectively suggest that the TM hydrophobicity may be an important determinant for Spc1-regulated SPase processing of membrane proteins (Figs. 5D and E).

### SPase-mediated cleavage of TM segments is decreased upon overexpression of Spc1

Since processing of test membrane proteins by SPase increased in the absence of Spc1, we wondered whether overexpression of Spc1 also affects processing, possibly in an opposite manner. The LepH2(5L) construct was radiolabeled and chased over 15 min in the WT strain containing an EV or Spc1 overexpression (OE) vector (Fig. 5F). Cleavage of LepH2(5L) in both strains was increased during the 15 min chase, indicating that cleavage continued post-translationally. At 0 min chase, cleavage of LepH2(5L) in the Spc1 OE cells was slightly reduced compared to that of in WT cells, but at 15 min chase, cleavage in Spc1 OE cells was markedly reduced compared to that of in WT cells (Figs. 5F and G). Under Spc1 OE conditions, cleavage of other test membrane proteins, double-spanning LepCC(17L) and single-spanning LepCCt(17L) variants also decreased compared to those expressed in WT cells (Figs. S4C and D). These data suggest that additional Spc1 can protect TM segments from SPase-mediated cleavage co- and post-translationally.

### Spc1 interacts with membrane proteins with uncleaved TM segments

How does overexpressed Spc1 protects TM segments from SPase-mediated processing? We suspected that Spc1 might physically shield TM segments from being presented to the SPase active site. If so, overexpressed Spc1 might interact with signal-anchored or membrane proteins, and we carried out co-immunoprecipitation to test the idea (Fig.6). The C-terminally FLAG-tagged Spc1 (Spc1FLAG) was overexpressed in the *spc1*Δ strain together with HA-tagged CPYt and LepCC model proteins. We assessed whether N9CPYt(*h*) Q-3/P+1’ that is membrane-anchored (Fig. S3D) interacts with Spc1, along with N9CPYt(*l*) and N9CPYt(*h*) variants carrying cleavable SSs. N9CPYt(*h*) Q-3/P+1’ was co-immunoprecipitated with Spc1 while N9CPYt(*l*) and N9CPYt(*h*) were not (Fig. 6A). Next, we tested LepCC(14L) and LepCC(17L) variants. Although LepCC(14L) variant was ∼70% cleaved in the WT strain (Fig. S4B), its cleavage was reduced less than 50% in the *spc1*Δ strain with Spc1 overexpression (Fig. 6B). While a cleaved product was not co-immunoprecipitated with Spc1, a full-length protein was (Fig. 6B). A full-length LepCC(17L) was also co-precipitated with Spc1 (Fig. 6B). These data show that Spc1 associates with membrane proteins having uncleaved TM segments, suggesting that Spc1 interacts with TM segments of membrane proteins and shield them from being processed by SPase.

**Figure 6.**
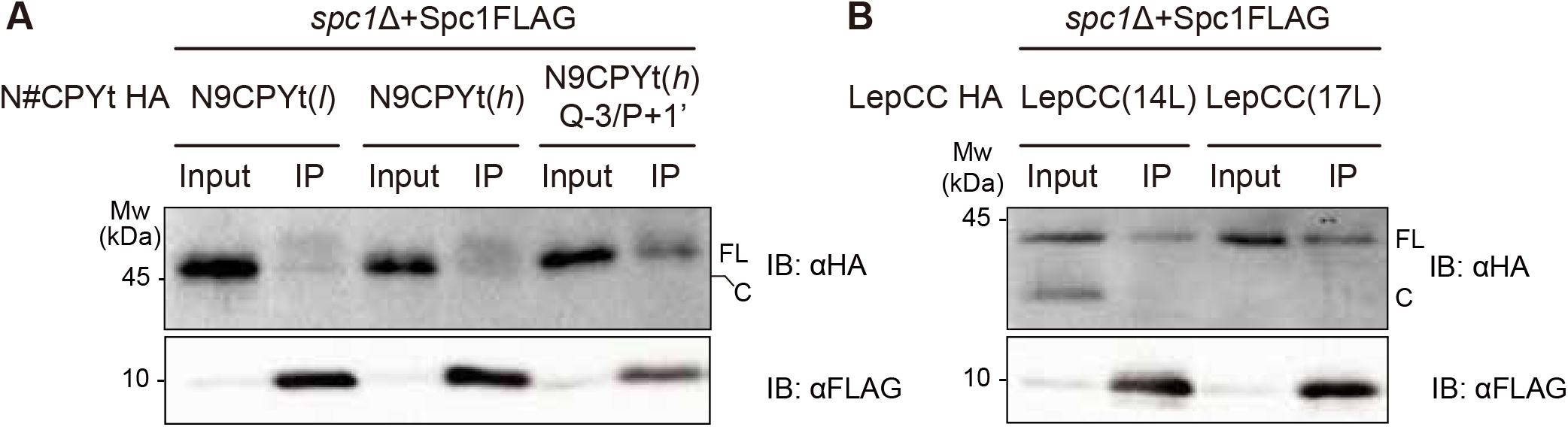
Overexpressed Spc1 interacts with uncleaved model proteins. Co-immunoprecipitation of overexpressed Spc1 and (A) N9CPYt and (B) LepCC variants. spc1Δ strain co-expressing FLAG-tagged Spc1 and HA-tagged indicated substrates was subjected to crude membrane fractionation. Isolated membrane was solubilized with lysis buffer containing 1% Triton X-100 and the resulting lysate was subjected to co-immunoprecipitation against FLAG (Spc1FLAG) via protein G-agarose and anti-FLAG antibody, visualized by SDS-PAGE and Immunoblotting (IB) with anti-HA antibody. IP; immunoprecipitation; FL, full length; C, cleaved.

## Discussion

The sequence context of SPs that are cleaved by SPase and signal-anchored segments that become TM domains are similar in their hydrophobicity and overall length. Hence, both can be recognized by the signal recognition particle (SRP), act as an ER targeting signal and initiate protein translocation in the ER membrane. In a subsequent step, an SP is cleaved, whereas a TM segment evades processing by SPase and anchors in the membrane. Although the cleavage motif is needed, it is not the sole factor for whether SPase cleaves the signal sequence or not. It remains elusive how SPase discriminates TM segments from SPs.

Analyzing the cleavage of CPY-based SSs of systematically varied N-length and hydrophobicity, our data show that the substrate spectrum of SPase is defined by the N-length and hydrophobicity of SSs; SSs with shorter *n* regions and/or less hydrophobic *h* regions are better substrates for SPase. Consistent with our findings, it has been observed that some type II single-pass membrane proteins are processed by SPase when their *n* region is shortened in mammalian cells (Lemire et al., 1997; Lipp and Dobberstein, 1986; Roy et al., 1993; Schmid and Spiess, 1988).

When the cleavage of the same CPY variants was assessed in the absence of Spc1, cleavage of internal signal-anchored sequences was markedly enhanced, and the cleavage pattern was restored when *SPC1* was re-expressed. There are two possibilities for the expanded CPY substrate spectrum of SPase in the absence of Spc1: 1) SPase may cleave SSs in sites other than the canonical site or 2) SPase may cleave signal peptide-like sequences such as TM domains due to compromised capacity for substrate selection. We carried out mutational analysis of the SS cleavage sites of CPY variants to test the first possibility and found that SPase processes only the canonical SS cleavage site with or without Spc1, excluding the first possibility. For the second possibility, we tested the processing of membrane proteins in the *spc1*Δ strain. A pulse-chase experiment showed that cleavage of TM segments in model membrane proteins in *spc1*Δ cells was enhanced at early time points of metabolic labeling and further increased at subsequent chase times, indicating that processing continues after membrane insertion. We also observed that SPase-mediated cleavage of TM segments of model membrane proteins was reduced upon overexpression of Spc1. These results suggest that the expanded CPY and Lep membrane protein substrate spectrum of SPase without Spc1 is due to its compromised capability of sorting out TM segments from SPase action.

How does Spc1 sort out TM segments from SPase? Our co-immunoprecipitation data show that Spc1 interacts with membrane proteins with uncleaved TM segments. These results suggest that Spc1 may shield TM segments from being presented to the active site of SPase, thereby protecting them. In turn, this function of Spc1 contributes to accurate substrate selection for SPase.

While Spc1 is dispensable for growth in yeast, deletion of SPC12 (Spc1 homolog) in *Drosophila* exhibits a developmental lethal phenotype (Haase Gilbert et al., 2013), suggesting that its function may be more prominent in higher eukaryotes. Intriguing observations have been made in SPCS1 knockout human cell lines. A genetic screen identified SPCS1 as one of the key regulators for the expression of ULBP1, a surface protein ligand for natural killer cells (Gowen et al., 2015). Genome-wide CRISPR screening identified SPCS1 as a key host factor in the processing of viral proteins that are made as polyproteins containing internal SSs and TM segments for the flavivirus family (Zhang et al., 2016). Interestingly, host SPCS1 was found to interact with the TM domains of viral proteins in Japanese encephalitis virus (Ma et al., 2018) and with the TM domains of viral proteins in hepatitis C virus (Suzuki et al., 2013). These observations indicate the involvement of SPCS1 in the regulation of processing and handling TM segments in higher eukaryotes.

Our study provides the evidence that SPase distinguishes SPs from TM segments and that Spc1 is involved in deselecting TM segments, thereby ensuring accurate substrate selection for SPase.

## Materials and Methods

### Yeast strains

The *S. cerevisiae* haploid W303-1α (*MATα, ade2, can1, his3, leu2, trp1, ura3*) was used as a WT strain. The *SPC1* ORF in W303-1α was replaced with *HIS3* amplified from the pCgH plasmid (Kitada et al., 1995) by homologous recombination to generate the *spc1*Δ strain (*MATα, spc1Δ::HIS3, ade2, can1, his3, leu2, trp1, ura3*). *spc3-4* is a temperature-sensitive mutant exhibiting a defect in SPase activity at 37°C (Fang et al., 1997). For the overexpression of Spc1, the pRS426 vector carrying *SPC1* under the GPD promoter was transformed into the W303-1α strain.

### Construction of plasmids

All CPY variants were derived from pRS424GPD N26CPY-HA constructed in our previous study (Yim et al., 2018). Using this construct as a template, we first truncated residues 323-532 of CPY by site-directed mutagenesis following the manufacturer’s protocol (Toyobo, Japan). Next, the N-terminus was truncated, and the hydrophobicity of the CPY SS was modified by site-directed mutagenesis. *E. coli* Lep-derived LepCC constructs (Nilsson et al., 1994) were subcloned from the pGEM4z vector into the yeast pRS424 vector by PCR amplification and homologous recombination. LepCCt constructs were generated by truncation of the N-terminal 20 residues except for the start methionine in pRS424GPD LepCC constructs. The pRS426 vector containing *SPC1* was cloned by homologous recombination or using the Gibson assembly kit following the manufacturer’s protocol. All plasmids were confirmed by DNA sequencing. LepH2 variants in a yeast vector were constructed in (Lundin et al., 2008).

### Pulse and pulse-chase experiments

Pulse labeling and pulse-chase procedures were carried out as in (Reithinger et al., 2014; Yim et al., 2018). Briefly, yeast cells were grown at 24-30°C until the OD_600_ reached between 0.3 and 0.8 in selective medium. Then, 1.5 OD_600_ units of cells were harvested by centrifugation (2,170×*g*, 5 min, 4°C), washed with -Met medium without ammonium sulfate, and incubated at 30°C for 10 min. Cells were centrifuged and resuspended in 150 μl of -Met medium without ammonium sulfate, and radiolabeled with [^35^S]-Met (40 μCi per 1.5 OD_600_ units of cells) for 5 min at 30°C. After incubation, labeling was stopped by the addition of 750 μl of ice-cold stop solution buffer containing 20 mM Tris-HCl (pH 7.5) and 20 mM sodium azide. Cell pellets were harvested by centrifugation (16,000×*g*,1 min, 4°C) and stored at - 20°C until use.

For pulse-chase experiments, 1.5 OD_600_ units of cells were harvested for each time point. Cells were prepared the same way as in pulse labeling, except that cells were resuspended in -Met medium of twice or three times the volume corresponding to the number of time points for chase. Radiolabeling was stopped and chased by the addition of 50 μl of 200 mM cold Met medium per 1.5 OD_600_ units of cells for each time point. The reaction was stopped by transferring 1.5 OD_600_ units of cells to 750 μl of ice-cold stop solution buffer and centrifuged, and the cell pellets were kept frozen at −20°C until use.

### Tunicamycin treatment

Tunicamycin treatment of growing cells for radiolabeling and autoradiography was carried out as described in (Yim et al., 2018). Briefly, prior to radiolabeling, cells were starved with 1 ml of -Met medium without ammonium sulfate for 30 min at 30°C in the presence of 100 μg/ml tunicamycin (Sigma) dissolved in DMSO, while control cells were mock-treated with DMSO.

### Immunoprecipitation and SDS-PAGE

Radiolabeled cell pellets were resuspended in 100 μl of lysis buffer (20 mM Tris-HCl (pH 7.5), 1% SDS, 1 mM DTT, 1 mM PMSF, and 1X Protease Inhibitor Cocktail) and mixed with 100 μl of ice-cold glass beads. Cell suspensions were vortexed for 2 min twice, keeping the samples on ice for 1 min in-between. Subsequently, the samples were incubated at 60°C for 15 min and centrifuged (6,000×*g*, 1 min, 4°C). Supernatant fractions were mixed with 500 μl of immunoprecipitation buffer (15 mM Tris-HCl (pH 7.5), 0.1% SDS, 1% Triton X-100, and 150 mM NaCl), 1 μl of anti-HA antibody, and 20 μl of prewashed protein G-agarose beads (Thermo Scientific Pierce) and rotated at room temperature for 3 h. The agarose beads were washed twice with immunoprecipitation buffer, once with ConA buffer (500 mM NaCl, 20 mM Tris-HCl (pH 7.5), and 1% Triton X-100), and once with buffer C (50 mM NaCl and 10 mM Tris-HCl (pH 7.5)). The beads were incubated with 50 μl of SDS sample buffer (50 mM DTT, 50 mM Tris-HCl (pH 7.6), 5% SDS, 5% glycerol, 50 mM EDTA, 1 mM PMSF, 1X Protease Inhibitor Cocktail (Quartett), and bromophenol blue) at 60°C for 15 min, followed by Endo H (Promega) treatment at 37°C for 1 h. Protein samples were then loaded onto SDS-PAGE gels and separated by electrophoresis.

### Data quantification

A Typhoon FLA 7000 phosphoimager and a Typhoon FLA 9500 phosphoimager were used for the detection of radiolabeled signals on SDS-PAGE gels by autoradiography. Data were processed and quantified using MultiGauge version 3.0 software. Cleavage efficiency was calculated from the band intensities of glycosylated bands using the formula: cleavage (%) = cleaved band × 100/cleaved + full-length bands. Cell growth assay and carbonate extraction samples were detected by ChemiDoc™ XRS+, and resulting data were processed using Image Lab™ software.

### Statistical analysis

Statistical analyses with obtained quantification data were performed using Microsoft Excel 2013 or GraphPad Prism 8 for Windows.

### Carbonate extraction

Carbonate extraction was carried out as described in (Yim et al., 2018) with the following modifications. Five OD_600_ units of cells were harvested by centrifugation (2,170×*g*, 5 min, 4°C), washed with dH_2_O and subjected to lysis. The final lysate was subjected to centrifugation (20,000×*g*, 30 s, 4°C) to remove cell debris and transferred to a new prechilled tube. Centrifugation (20,*000×g*, 20 min, 4°C) followed after incubation with 0.1 M Na_2_CO_3_ (pH 11.5). Trichloroacetic acid (TCA)-precipitated ‘Total’, ‘Supernatant’ and ‘Pellet’ fractions were centrifuged (20,*000×g*, 15 min, 4°C) and washed with acetone. Samples were resuspended in SDS sample buffer and analyzed by SDS-PAGE and Western blotting.

### *In vitro* transcription/translation

For *in vitro* protein synthesis, the TnT Quick Coupled SP6 Transcription/Translation System (Promega) was used by following the manufacturer’s protocol. The pGEM-4Z plasmid containing ΔSS-CPY (signal sequence of CPY deleted) was incubated with TnT SP6 Quick Master Mix and [^35^S]-Met for 1 h at 30°C. The synthesized proteins were then analyzed by SDS-PAGE and autoradiography.

### Mass spectrometry analysis of the abundance of SPC subunits in WT and *spc1*Δ strains

Cells from the W303-1α and *spc1*Δ strains were grown overnight in YPD medium at 30°C in biological triplicates. Fifteen OD_600_ units of cells were harvested for each strain and subjected to cell lysis by vortexing for 10 min at 4°C with 200 μl of lysis buffer (8 M urea, 1x PIC, and 1 mM PMSF) and glass beads. The resulting cell lysates were reduced and alkylated with 10 mM dithiothreitol and 40 mM iodoacetamide, respectively, followed by trypsinization after 10-fold dilution with 50 mM ammonium bicarbonate buffer. The digested samples were then subjected to further clean-up with a C18 cartridge.

For quantitative analysis, 10 μg of the biological triplicate of individual samples was subjected to TMT labeling as follows: TMT-126, 128N, and 130C for W303-1α samples; TMT-127C, 129N, and 130N for *spc1*Δ samples. The TMT-labeled peptide sample was subjected to LC-MS3 analysis using Orbitrap Fusion Lumos (Thermo Fisher Scientific) with the following mass spectrometric parameters. The 10 most intense ions were first isolated at 0.5 Th precursor isolation width under identical full MS scan settings for CID MS2 in an ion trap (ITmax 150 ms and AGC 4E3). The 10 most intense MS2 fragment ions were synchronously isolated for HCD MS3 (AGC 1.5E5, ITmax 250 ms, and NCE 55%) at an isolation width of 2 *m*/*z*.

### Co-immunoprecipitation

Experimental procedures are based on (Zhang et al., 2017) with following modifications. Crude membrane was isolated from about 15 OD_600_ units of cells and solubilized with 400 μl of lysis buffer (50 mM HEPES-KOH/PBS, pH 6.8, 1% Triton X-100, 150 mM KOAc, 2 mM Mg(OAc)_2_, 1mM CaCl_2_, 15% glycerol, 1x PIC, 2 mM PMSF) by rotation at 4°C for 1h. After centrifugation at 14800 rpm for 30 min at 4°C, soluble fraction was transferred to a tube containing 25 μl of protein G-agraose beads prewashed three times with lysis buffer, followed by rotation for 30 min at 4°C. Beads were removed by quick centrifugation and 15 μl of the lysate was saved for ‘Input’ fraction while all the remaining supernatant was transferred to a new tube containing 25 μl of prewashed protein G-agraose beads and 1 μl of anti-FLAG mouse antibody (FUJIFILM Wako Pure Chemical Corporation). The immunoprecipitation mixture was rotated for about 4h at 4°C. Beads were washed three times with lysis buffer and sampled by incubation with 40 μl of SDS sample buffer for 15 min at 55°C as ‘IP’ fraction. ‘Input’ fraction was mixed with 65 μl of SDS sample buffer and incubated for 15 min at 55°C.

## Acknowledgments

Authors thank Professor Gunnar von Heijne (Stockholm University, Sweden) for kindly providing *E*.*coli* Lep constructs and reading of the manuscript, and the lab members for discussion and critical reading of the manuscript.

## Author Contributions

C. Y., Y. C. and H. K. designed research; C. Y. and Y. C. performed experiments using yeast cells and analyzed data; J. K. and J-S. K performed and analyzed mass spectrometry data; I. N. provided Lep constructs; C. Y., Y. C. and H. K. wrote the paper; C. Y., Y. C., J. K., I. N., J-S. K, and H. K. read and edited the manuscript.

---

**Figure 1-figure supplement 1.**
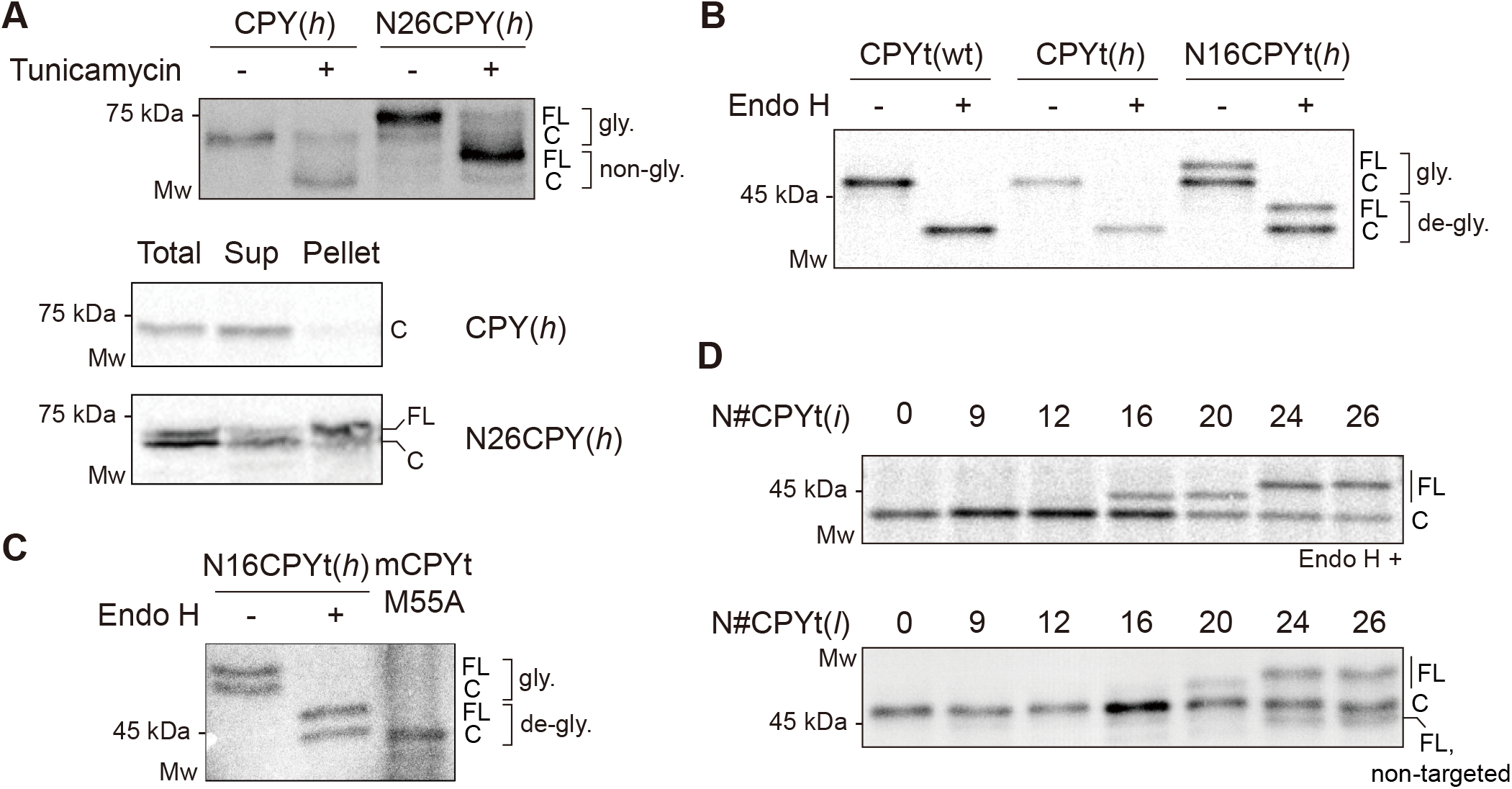
SPase-mediated processing of CPY variants depends on the N-length and hydrophobicity of the SS. (A) *Top*, CPY(*h*) and N26CPY(*h*) constructs in WT cells were radiolabeled for 5 min at 30°C in the presence or absence of tunicamycin, followed by immunoprecipitation and SDS-PAGE and analysis by autoradiography. *Bottom*, carbonate extraction was carried out. Sup, supernatant. (B) CPYt(wt), CPYt(*h*), and N16CPYt(*h*) constructs in WT cells were radiolabeled for 5 min at 30°C and subjected to immunoprecipitation for protein sampling. All protein samples were treated with Endo H prior to SDS-PAGE and analyzed by autoradiography. (C) N16CPYt(*h*) in the WT strain was radiolabeled for 5 min at 30°C, and the resulting protein sample was compared with the *in vitro* translated SS-deleted mature CPY (mCPYt M55A) on an SDS-PAGE gel. mCPYt M55A, in which M55 was substituted to alanine to silent the alternative start codon. (D) N#CPYt(*i*) (top) and N#CPYt(*l*) (bottom) variants in WT cells were analyzed as in Fig. 1B. In the N#CPYt(*l*) variants, a minor amount of the unglycosylated band was detected when the N-length became longer than 24, indicating inefficient translocation.

**Figure 2-figure supplement 1.**
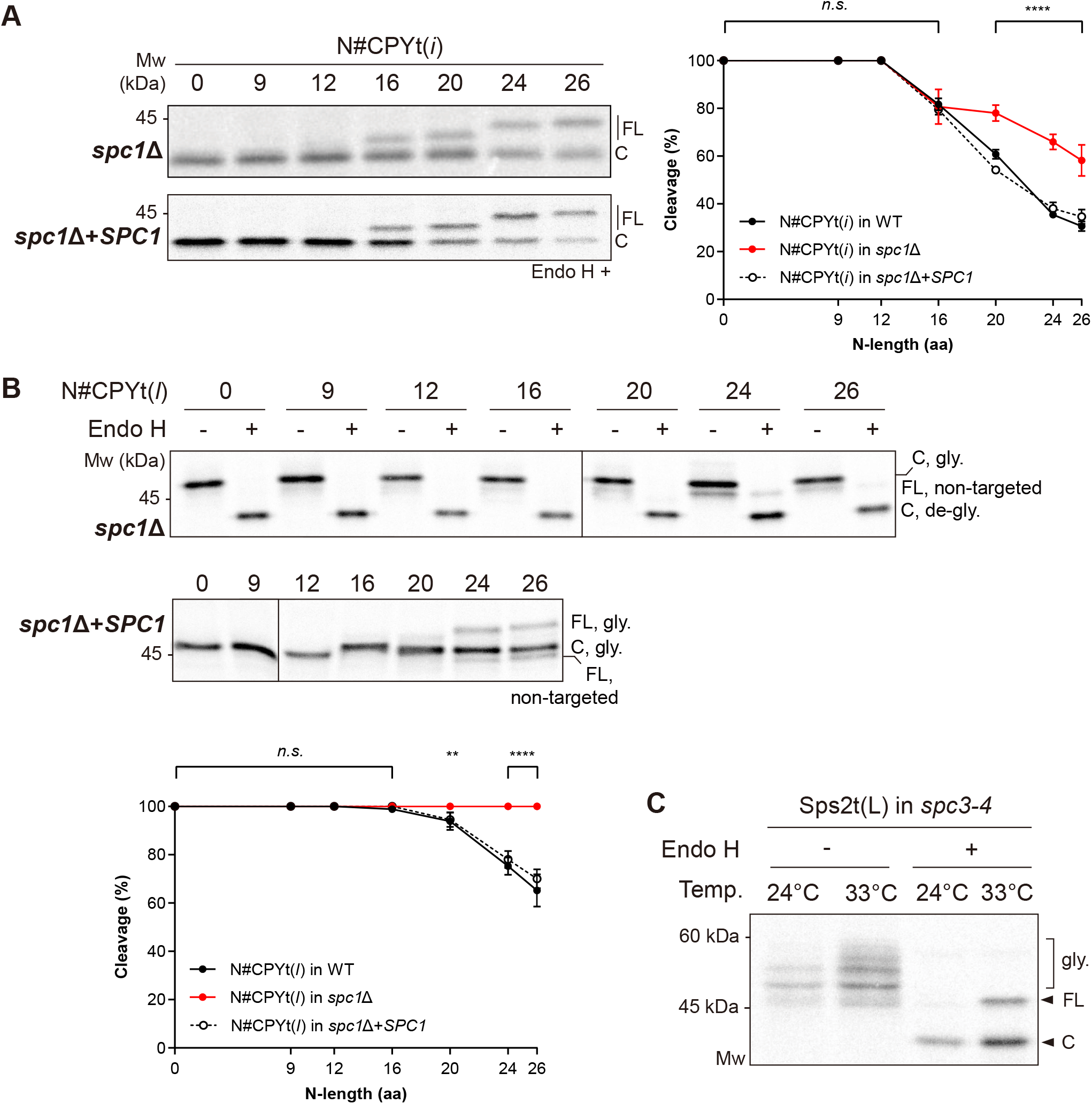
Cleavage of internal SSs is increased in the absence of Spc1. (A) N#CPYt(*i*) variants in *spc1*Δ and *spc1*Δ+*SPC1* cells were expressed, and protein samples were prepared as in Fig. 1B. Cleavage was analyzed as in Fig. 1F, and the data from the WT, *spc1*Δ and *spc1*Δ+*SPC1* strains were compared. (B) N#CPYt(*l*) variants in the WT, *spc1*Δ and *spc1*Δ+*SPC1* strains were analyzed, and the data were compared as in (A). p-values were calculated by multiple t-tests; *n*.*s*., p>0.05; **, p≤0.01; ****, p≤0.0001. (C) Processing of Sps2t variants in the *spc3-4* strain at 24°C or 33°C.

**Figure 3-figure supplement 1.**
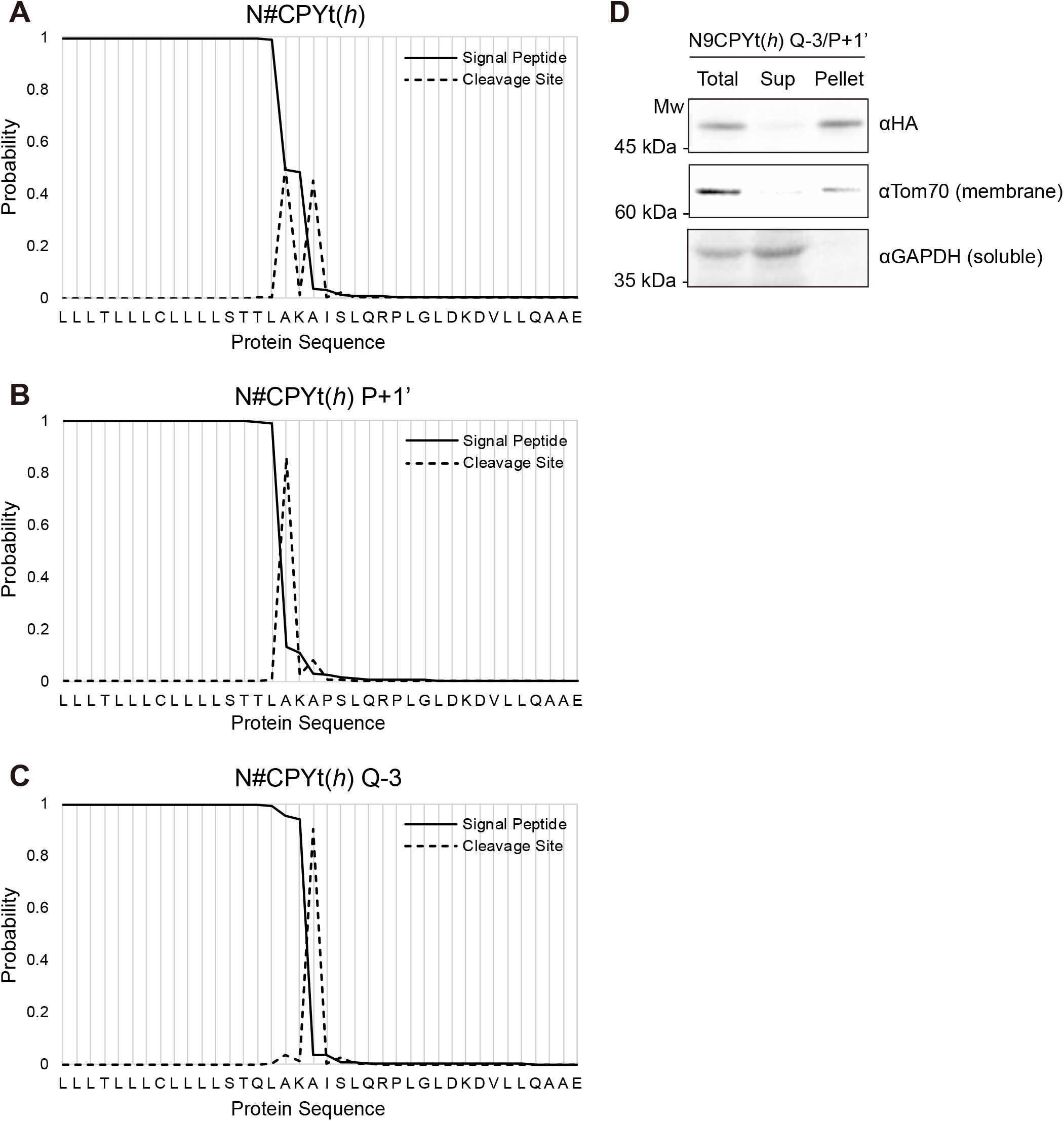
Prediction of cleavage sites in CPY SSs. Cleavage sites were predicted using SignalP 5.0 software (http://www.cbs.dtu.dk/services/SignalP/) (Almagro Armenteros et al., 2019). (A) N#CPY(*h*), (B) N#CPY(*h*) P+1’, (C) N#CPY(*h*) Q-3. Peaks of the dashed line indicate the predicted cleavage sites. (D) Carbonate extraction of N9CPYt(*h*) Q-3/ P+1’. Sup, supernatant fraction. Anti-Tom70 and anti-GAPDH antibodies were used as controls for membrane and soluble proteins, respectively.

**Figure 5-figure supplement 1.**
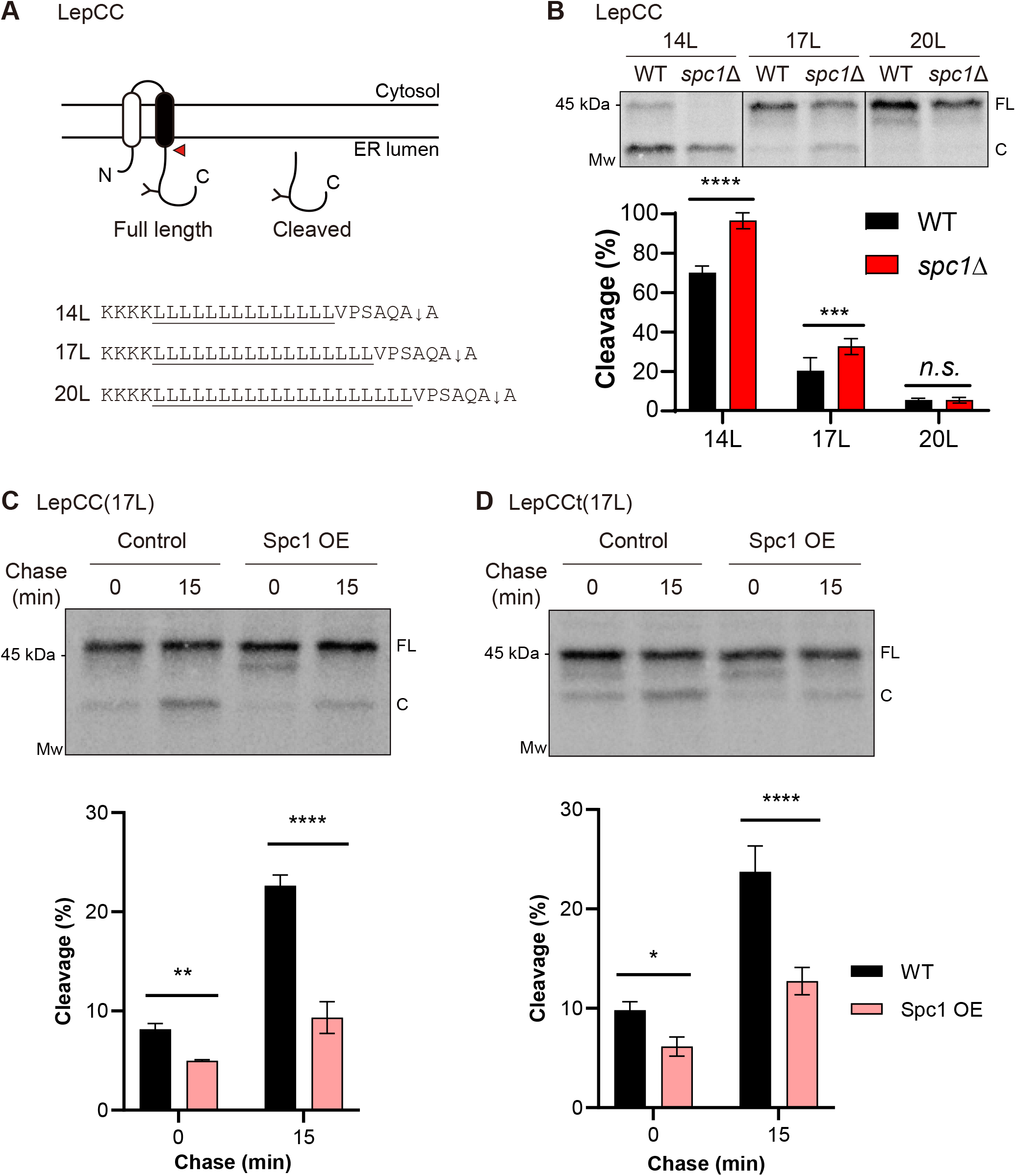
SPase-mediated processing of membrane proteins is modulated by Spc1. (A) Schematics of LepCC. The H segment is colored black, and an N-linked glycosylation site is indicated as Y. Flanking and TM sequences of an H segment are shown for three LepCC variants. The cleavage site is shown as an arrow (↓). A red arrowhead points to cleavage by SPase. (B) LepCC variants in the WT or *spc1*Δ strain radiolabeled for 5 min at 30°C were analyzed as shown in Fig. 4. At least three independent experiments were carried out. The representative blot is shown (*top*) and the average is shown with the standard deviation (*bottom*). (C and D) LepCC(17L) (C) or LepCCt(17L) (D) in WT cells harboring control vector or Spc1 overexpression (OE) vector. Transformants were subjected to radiolabeling for 5 min at 30°C followed by chasing for the indicated time points. Cleavage (%) was calculated as in Fig. 1F. For all the experimental sets, three independent experiments were carried out, and the average is shown with the standard deviation. FL, full length; C, cleaved. p-values were calculated by multiple t-tests; *n*.*s*., p>0.05; *, p≤0.05; **, p≤0.01; ***, p≤0.001; ****, p≤0.0001.

